# Simulating individually targeted transcranial electric stimulation for experimental application

**DOI:** 10.1101/739904

**Authors:** Jan-Ole Radecke, Asad Khan, Andreas K. Engel, Carsten H. Wolters, Till R. Schneider

**Author notes:** Corresponding author at: Department of Neurophysiology and Pathophysiology, University Medical Center Hamburg-Eppendorf, Martinistr. 52, 20251 Hamburg, Germany. Declarations of interest: None. Funding: This work was supported by the German Research Foundation (DFG) [SPP 1665/SCHN 1511/1-2 to TRS; WO1425/5-2 to CHW].

## Abstract

Transcranial electric stimulation (tES) induces electric fields that are subject to a complex interaction with individual anatomical properties, such as the low-conducting human skull, the distribution of cerebrospinal fluid or the sulcal depth, as well as stimulation target location and orientation. This complex interaction might contribute to the heterogenous results that are commonly observed in applications of tES in humans. Targeted tES, on the other hand, might be able to account for some of these individual factors. In the present study, we used the finite-element method (FEM) and head models of twenty-one participants to evaluate the effect of individually targeted tES on simulated intracranial current densities. Head models were based on an automated segmentation algorithm to facilitate processing in experimental sample sizes. We compared a standard stimulation montage to two individually optimized tES montages using an Alternating Direction Method of Multipliers (ADMM) and a Constrained Maximum Intensity (CMI) approach. A right parietal target was defined with three different orientations. Individual current densities showed varying intensity and spatial extent near the lower limit at which physiological efficacy of electric fields can be assumed. Both individually optimized targeting algorithms were able to control the electric field properties, with respect to intensities and/or spatial extent of the electric fields. Still, across head models, intensity in the stimulation target was constrained by individual anatomical properties. Thus, our results underline the importance of targeted tES in enhancing the effectiveness of future tES applications and in elucidating the underlying mechanisms.

## 1. Introduction

*Transcranial electric stimulation* (tES) is commonly used as a non-invasive tool to modulate neuronal activity in healthy and clinical samples (Lefaucheur, 2016; Nitsche et al., 2008; Nitsche and Paulus, 2011; Schutter and Wischnewski, 2016). However, a large variability regarding effects of tES on behavior and neurophysiology was observed (López-Alonso et al., 2014; Schouwenburg et al., 2018; Veniero et al., 2017, 2015; Wiethoff et al., 2014), hampering the understanding of determinants underlying tES (Bestmann et al., 2015; Horvath et al., 2015a, 2015b). Besides the endogenous physiological properties of the stimulation target (e.g. Thut et al., 2017), the strength and spatial distribution of the induced electric field within the brain is affecting the tES efficacy (Kasten et al., 2019; Laakso et al., 2015; Opitz et al., 2015). Recently, it was suggested to include simulations of the induced electric field in tES studies as a standard or even adapt the stimulation to the individual brain a priori (Berker et al., 2013; Bestmann et al., 2015; Liu et al., 2018; Polanía et al., 2018).

In typical tES applications a spatially specific brain area is targeted by the electric field of the stimulation. Besides the variable localization of perceptual and cognitive functions, individual anatomy shows a large variability, affecting the estimated electric field with respect to its magnitude and orientation in the cortex (Dmochowski et al., 2013; Huang et al., 2017; Laakso et al., 2015; Opitz et al., 2015; Truong et al., 2013; Wagner et al., 2014). Therefore, the simulation of intensity and orientation of the electric field in the individual brain is of great importance to guide stimulation of the target region in a physiologically efficient way (Creutzfeldt et al., 1962; Krieg et al., 2015, 2013; Nitsche and Paulus, 2000; Radman et al., 2009). The target orientation plays an important role, since the electric field exerts its effect along the field orientation but has no effect for vectors orthogonal to the field orientation (Krieg et al., 2015, 2013). Cortical pyramidal cells show a systematic alignment of their apical dendrites radial to the cortical surface. Therefore, assemblies of pyramidal cells can be modeled as a summed dipole and might be especially susceptible to changes in the orientation of the electric field (Creutzfeldt et al., 1962; Krieg et al., 2015, 2013; Nitsche and Paulus, 2000; Radman et al., 2009).

In healthy human subjects, the scope of applicable methods for the evaluation of current flow within the brain is restricted to non-invasive approaches. The finite-element method (FEM) is widely used to model the complex volume conductor for the simulation of tES-induced current flow inside the brain (Datta et al., 2009; Wagner et al., 2014, 2007; Windhoff et al., 2013). FEM models of the human head are typically derived from MRI scans by segmenting the head volume into different tissue compartments with specific conductivities assigned to each tissue type. Based on the estimation of electric fields in the brain, the electrode placement and the applied current strength can be inversely defined to optimally target a specified brain region or dipole within the brain (Dmochowski et al., 2011, 2017; Guler et al., 2016; Ruffini et al., 2014; Wagner et al., 2016a). Using individual FEM head models, it is thus possible to adapt the stimulation montage to intra- and inter-individual anatomical properties (Dmochowski et al., 2013; Opitz et al., 2015). Stimulation intensity and focality of the electric field, which are determined by the stimulation parameters, are intended to be optimized for the specified target. In principle, a compromise between these two properties of the stimulation has to be reached in every montage, yielding either a higher intensity with lesser focality or vice versa (Dmochowski et al., 2011). Therefore, the algorithms to optimize the stimulation montage either aim at improving the intensity or constrain the spatial extent of the electric field with respect to a specified intracranial target (Dmochowski et al., 2011; Guler et al., 2016; Wagner et al., 2016a).

However, FEM modeling has not yet been widely used in experimental studies with larger sample sizes (Boayue et al., 2018; Kasten et al., 2019; Laakso et al., 2015), as the construction of FEM head models is computationally demanding and the segmentation of the anatomy is still often performed manually or semi-automatically. Furthermore, individually targeted tES has only been described occasionally, for example in a small sample of stroke patients (Dmochowski et al., 2013). Other groups defined the stimulation montage based on the topographical EEG voltage distribution in patients with Parkinson’s disease (Del Felice et al., 2019). This approach was previously termed *naive reciprocity*, due to volume conduction effects that are neglected during the inverse definition of the stimulation montage based on EEG sensor data (Dmochowski et al., 2017). One previous study used various initial stimulation montages, picking the one montage that showed the highest impact on a verbal task for a subsequent aphasia training (Shah-basak et al., 2015). To our knowledge, no study thus far evaluated targeted electric fields, based on algorithmic optimization in a sample size that is able to reliably capture individual variability across subjects.

In contrast, in the large majority of tES studies a standardized stimulation montage is applied to all participants, neglecting differences in individual anatomy, as well as target location and orientation. Despite the undisputed merit of such studies, it is reasonable to assume that applying the same standard montage to different individual anatomies would lead to a high chance of misdirecting the electric field with respect to target intensity and orientation. Consequently, functionally different neural assemblies in each participant are stimulated that would result in varying effects on behavioral or neurophysiological outcome measures. Therefore, it is important to understand the relationship between individual brain structure and the high variability in the efficacy of tES applications. Previous publications support the notion that electric field distributions have an impact on the actual efficacy of tES (Edwards et al., 2013; Kasten et al., 2019; Mendonca et al., 2011). In order to control the influence of anatomical variation, the stimulation montages can be optimized for individual stimulation targets and anatomy (Dmochowski et al., 2011, 2017; Guler et al., 2016; Ruffini et al., 2014; Wagner et al., 2016a).

In the present study, we simulated tES-induced electric fields in FEM-head models of twenty-one participants. We compared three different methods to determine the stimulation montages from which the first two were individualized: The *Constrained Maximum Intensity (CMI*) algorithm (Khan et al., 2019) and the focality-optimizing *Alternating Direction Method of Multipliers (ADMM)* algorithm (Wagner et al., 2016a). The third method was a standard *5×1* montage, which was not individually optimized. Stimulation targets with three different orientations were defined, based on a standard brain atlas. Target 1 was radial to the scalp surface (*radial, RAD*), target 2 was tangential to the scalp surface with an anterior-posterior orientation (*tangential_a-p_, TAP*), and target 3 was tangential with a left-right orientation (*tangential_l-r_, TLR*). This approach allows to determine the stimulation parameters with respect to individual structure, as well as target location and orientation, to evaluate the electric fields that are induced by individually targeted tES and to compare between the three methods. To assess the variability of parameters over subjects, realistic FEM head models were computed for a sample with the size of typical tES experiments.

We hypothesize that targeted tES montages computed with the CMI and ADMM algorithms will surpass the standard montage with respect to target field strength and spatial focality, respectively. Furthermore, we assume that targeted tES is able to address the variability of the electric fields that is introduced by individual brain structure and different target orientations.

## 2. Materials and Methods

Optimization of stimulation electrode montages was applied, based on individual and automatically segmented *six compartment FEM (6C FEM)* head models using MATLAB (Natick, MA, USA), SPM12 (www.fil.ion.ucl.ac.uk/spm/), FieldTrip (Oostenveld et al., 2011), and METH (Nolte) toolboxes, as well as custom MATLAB-scripts. Forward solutions were computed with the open-source toolbox SimBio (SimBio Development Group).

### 2.1 Participants

*Twenty-one* right-handed participants (12 female, 28.2 ± 4.7 years [range 20 - 37]) were included in this study. All participants reported no history of neurological or psychiatric disorders and had normal or corrected-to-normal visual acuity and hearing. Participants were reimbursed for participation and gave written informed consent prior to the experiment. The experiment was conducted in agreement with the declaration of Helsinki and the protocol was approved by the ethics committee of the Hamburg Medical Association (Ärztekammer Hamburg).

### 2.2 MRI Data Acquisition and Automatic 6C Segmentation

For each subject structural T1 and T2-weighted (T1, T2) magnetic resonance (MR)-images were recorded with a 3T MR-scanner and a 64-channel head coil at an isotropic voxel resolution of 1×1×1 mm (Siemens Magnetom Prisma, Erlangen, Germany). Both, T1 and T2 images were acquired with an MP-RAGE pulse sequence (T1: TR/TE/TI/FA = 2300 ms/ 2.98 ms/ 1100 ms/ 9°, FoV = 192 x 256 x 256 mm; T2: TR/TE = 3200 ms/ 408 ms, FoV = 192 x 256 x 256 mm).

Individual, isotropic and geometry-adapted hexahedral FEM head models were computed (Aydin et al., 2014; Wagner et al., 2016b, 2014; Wolters et al., 2007), based on a new automatic six compartment (6C) segmentation algorithm. These head models were utilized for the simulation of electric fields induced by tES.

Structural T1 images were non-linearly co-registered onto the T2 images. Both T1 and T2 images were then separately segmented into five compartments using SPM 12 (white matter, gray matter, bone, skin, cerebrospinal fluid) (Ashburner and Friston, 2005). The resulting probability maps were further processed to produce *gray matter* (GM), *white matter* (WM), *cerebrospinal fluid* (CSF), *skin* (SKIN), *bone compacta* (BONE_C_) and *bone spongiosa* (BONE_S_) compartments (Fig. 1A-D).

**Fig 1.**
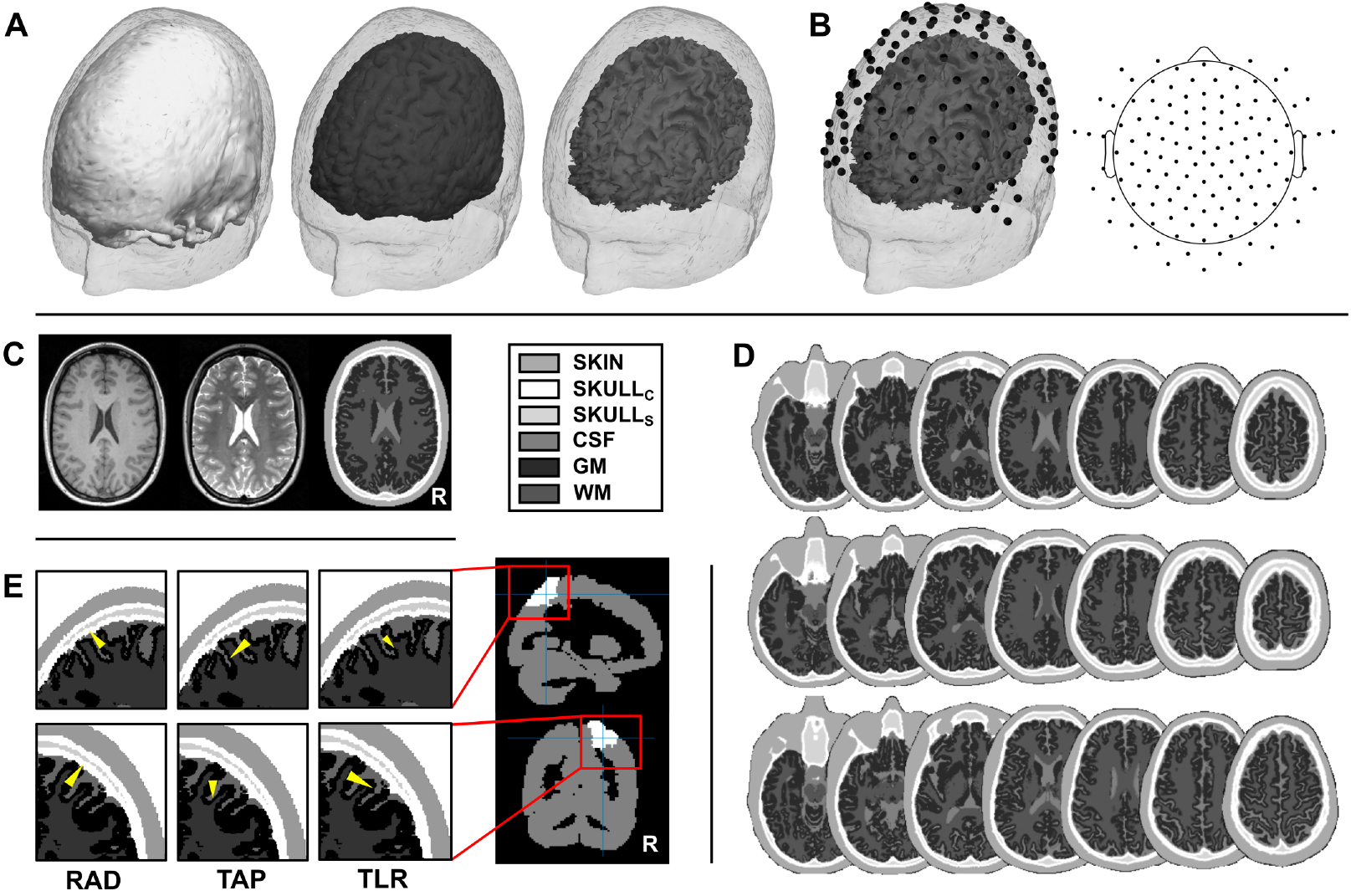
A) Solid isosurfaces of BONE_C_ (left), GM (middle) and WM (right) compartments, plotted within the transparent SKIN compartment for one exemplary participant. CSF and BONE_S_ where not displayed as isosurfaces, due to their disjointed structure. B) Simulated electrodes projected to the skin surface for the same example subject and shown with respect to WM (left) and as topographical illustration of the 126-channel layout (right). C) Correspondence between T1 and T2 MR images with the 6C segmentation in an axial slice of the same participant shown in A) and B). D) Three exemplary MRI segmentations show the reliability of the applied automated segmentation algorithm in axial slices. E) Stimulation target orientations (yellow cones) plotted on the 6C segmented volume as a close-up of the target region of interest in right SPL. Right SPL is highlighted (white) on the overall volume that is represented by the AAL atlas. Compartments are labeled as BONE_C_: bone compacta, BONE_S_: bone spongiosa, CSF: cerebrospinal fluid, GM: gray matter, WM: white matter.

In parallel to the initial SPM 12 segmentation, a binary head mask was estimated from the joint information of T1 and T2 scalp segmentations drawn from FieldTrip. Imaging artifacts outside the head volume were rejected. Ambiguous values from the initial SPM 12 segmentation were eliminated, by thresholding the probability maps for each probability map separately. T1 images provide a proper separation of gray and white matter. Thus, only corresponding T1 probability maps were further processed for GM and WM compartments. T2 imaging provides a better representation of fluids, compared to T1, therefore initial CSF estimates were taken from the T2 probability maps. A second bone probability map was constructed by averaging the initial T1 and T2 probability maps. The inner and outer boundaries of the skin compartment were taken from the averaged T1 and T2 information as well. As an interim result, each of the five tissue types was represented by one probability map with continuous voxel-wise probability values.

For each voxel, the largest probability values across tissue types defined the label of this voxel as belonging to the respective compartment. A binary brain mask was created including CSF, GM and WM compartments. Bone and skin segmentation artifacts included by this mask were eliminated. A binary bone mask was constructed from the integrated bone probability map, and adjacent to the brain mask. To dissociate BONE_C_ and BONE_S_, the binary bone mask was eroded and thresholded, based on the original T2 probability maps (Aydin et al., 2014; Wagner et al., 2014). The final SKIN compartment was estimated by the head mask, defining the outer boundary and the combined bone and brain mask to define the inner boundary of the skin. Due to image processing, interpolation, and removal of segmentation artifacts along these processing steps, tissue labels were partly overlapping. In order to integrate information of the different binary compartment masks into one volume, ambiguous labels were removed, while fixating BONE_C_ and BONE_S_. This constraint is necessary to assure that BONE_S_ is surrounded by a layer of BONE_C_ and leakage artifacts during the modeling of current flow are alleviated (Engwer et al., 2017) or avoided (Vorwerk et al., 2017). Missing tissue labels were interpolated iteratively using the nominal information from neighboring voxel labels.

The head volume was transformed from the SPM (RAS) to the CTF (ALS) coordinate system, defining the fiducial points individually as the nasion and the bilateral tragi, in order to align electrode positions with the head models. The lower third of the volume was cut off to reduce the overall size of the head model. Finally, a geometry-adapted hexahedral finite-element mesh was computed for the 6C head model volume of each subject. Electrodes were simulated using a point electrode model (Pursiainen et al., 2018), by projecting each electrode position to the closest FEM node on the scalp.

### 2.3 Definition of Stimulation Channel Locations and Simulated Targets

Individual electrode positions from 126-channel EEG caps (EasyCap, Herrsching, Germany) were optically registered (Xensor, ANT Neuro, Hengelo, The Netherlands) and averaged to a realistic, but standardized template across all subjects. The template electrode positions were aligned to the individual head models, directed by the fiducial points (nasion and bilateral tragi, Fig. 1B).

In practice, tES with small electrodes (Datta et al., 2009) can be used to replace the standardized EEG electrodes in the cap for combined EEG-tES applications. The relatively high spatial sampling of electrodes allowed the optimization algorithms to flexibly adapt electrode positions to the respective stimulation target and head model. Furthermore, in practice the close distance of electrode positions enables the stimulation current to be split among a number of electrodes. Thus, still at increasing stimulation intensity, skin sensations can be minimized (i.e. electric charge per electrode), while preserving a minimal loss of focality of the electric field. The overall simulation current was scaled to 2 mA (i.e. 4 mA peak-to-peak). In order to compare the different tES methods, the number of stimulation electrodes was fixed to *N* = 6 for each individual stimulation montage.

Stimulation targets were defined in the right *superior parietal lobule* (SPL) as a region of interest, based on the AAL brain atlas (*Automated Anatomical Labeling*) (Tzourio-Mazoyer et al., 2002). The target coordinate was determined as the average coordinate of all grid points within the right SPL in MNI-space. Individual 3D-grids (5 mm), sampling the combined CSF, GM and WM compartments were defined for each participant. By linear normalization of the individual T1 images on the MNI152 template brain (Montreal Neurological Institute, Montreal, Canada), the individual grids were warped into standardized MNI space. The inverse of the transformation matrix was applied to the target right SPL coordinate to warp it into the individual CTF coordinate system. Target coordinates were projected on the closest GM node of the individual head models. Finally, three orthogonal target orientations were simulated with respect to the individual scalp surface (Fig. 1E). One orientation was defined as radial to the scalp surface. Two tangential target orientations were computed along the anterior-posterior axis and along the left-right axis, respectively. A non-target coordinate was defined in the contralateral left SPL to relate the target current densities to an unstimulated area. Non-target orientations were defined as radial, tangential_a-p_ and tangential_l-r_ with respect to the scalp surface, respectively.

### 2.4 Inverse Optimization of Stimulation Montages and Forward Modeling of Current Flow

We compared two individually targeted tES montages with a standard montage. In principal, current densities within the brain can be modeled for the distribution of each certain electric field that is applied to the head transcranially. By combining the individual *m* electrode positions and 6C FEM head models with *n* nodes (approx. 3.5 - 4 million), we computed the matrix *A*. The matrix holds the forward model vectors *a_n_(r_n_)* at *r*_*n*_, which is the *n*_th_ node of the FEM model, while assuming a fixed current *s*_*m*_ applied to every combination of a fixed reference electrode and the *m*_th_ electrode of the given 126-channel layout (Dmochowski et al., 2011). Due to Helmholtz’ reciprocity, the forward model in tES inverse problems is symmetric to the lead field in EEG inverse problems, which reflects the mapping of the current source vectors to the electrode currents (Dmochowski et al., 2017; Wagner et al., 2016b). The current density vector field *j* at each node *r*_*n*_ for a given stimulation montage is determined by the linear combination of the weighted current *s* (including the reference electrode) applied to any combination of the *M* electrodes as a function of *A*:

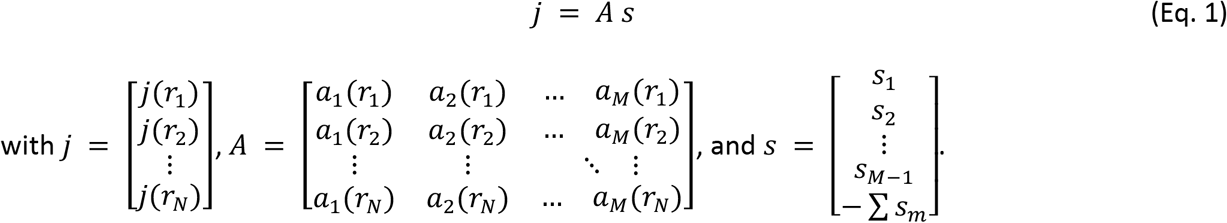

The tissue conductivities, which are input to the forward model vectors *a*_*n*_, were defined as 0.33 (GM), 0.14 (WM), 1.79 (CSF), 0.43 (SKIN), 0.025 (BONE_S_) and 0.007 (BONE_C_), as well as 1.4 S/m for the electrodes (Wagner et al., 2016a, 2014). The adjoint method (Wagner et al., 2016b) was employed to approximate the current densities induced by tES. Current densities were computed for all elements of the head models using the SimBio toolbox (SimBio Development Group). Previously, it was stated that the quasi-static approximation to Maxwell’s equations that was applied here, is not only applicable for direct currents, but also for low-frequency alternating current (< 1 kHz) (see Wagner et al., 2014). Consequently, the applied results should not only be valid for *transcranial direct current stimulation* (tDCS), but also for most applications of *transcranial alternating current stimulation* (tACS), or low-frequency *transcranial random noise stimulation* (tRNS).

The standard montage was defined as a 5×1 small electrode montage (cf. Datta et al., 2009), arranged around the center electrode that is placed closest to the radial projection of the target coordinate to the scalp. Due to the high-density 126-channel layout, five stimulation electrodes were placed around the center electrode to approximate a ring (comparable to 4×1 montages in a 64-channel layout). The distance between the center-electrode and surrounding electrodes was optimized manually, with respect to intensity and spatial extent of the stimulation.

The matrix *A* can be used to find the best possible weighting of current at the stimulation electrodes *S*_*max*_ under a defined set of constraints. The here applied optimization algorithms both compute the optimal stimulation montages either maximizing the target current density or optimizing the stimulation montage to balance stimulation intensity and focality.

#### 2.4.1 The CMI Algorithm

The originally proposed Maximum Intensity algorithm maximizes the current density along the target vector direction using an L1-norm which enforces the stimulation montage to produce a single anode and a single cathode (Dmochowski et al., 2011). In a subsequent formulation an additional maximum norm constraint was introduced, in order to limit the current applied to each electrode (Dmochowski et al., 2013). In the present study, we used a modified version of the Maximum Intensity algorithm, the *Constrained Maximum Intensity (CMI)* approach (Khan et al., 2019), which is expressed as:

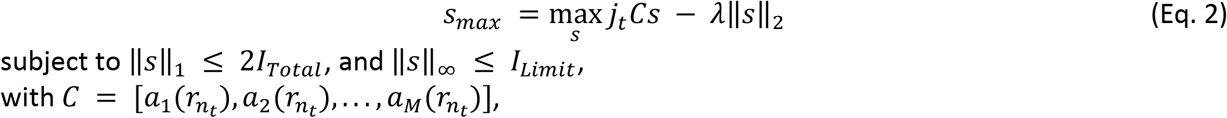

where *n*_*t*_ denotes the index of the target node and *j*_*t*_ the target current density vector. C is the submatrix from A which reflects the mapping of the electrode currents to the current density at the target vector. In this approach the maximum current applied to each electrode can be limited by a maximum norm constraint. Additionally, the applied current is distributed among stimulation electrodes by introducing the L2-norm (Khan et al., 2019). In our simulations, we chose λ = 250 to distribute the injected currents over the electrodes. *I*_*Total*_ = 2 mA was set to fulfill the safety constraint and *I*_*Limit*_ = 0.95 mA to enforce a distribution of electrode currents and also reduce the tactile perception of the stimulation under each electrode. In a two-step procedure, the stimulation montage was fixed to six electrodes for each individual stimulation montage.

#### 2.4.2 The ADMM Algorithm

The ADMM algorithm maximizes the current density in the stimulation target Ω_*t*_, while controlling the current densities in the remaining non-target volume conductor Ω. As described in detail by Wagner and colleagues (2016a), we considered an optimal control problem for a Laplace equation with Neumann boundary conditions with control and point-wise gradient state constraints. For numerical solution of the corresponding discretized problem the Alternating Direction Method of Multipliers (ADMM) was employed:

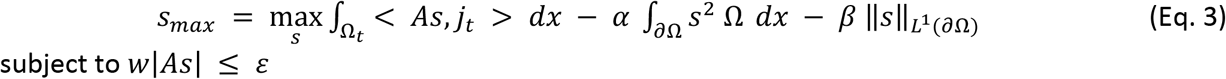

Herein, *w* reflects the weighting matrix that mediates between current densities in target and non-target regions. The parameter ϵ from the gradient state constraint reflects the upper bound of current densities in non-target regions and was set to 0.5 to enable focality of the solution. The L2-regularization parameter α penalizes the energy of the applied current pattern. The L1-regularization parameter β is used to minimize the number of active electrodes. We set both of these parameters to 0.0001 (elastic net) in order to guarantee a unique minimizer. A two-step procedure ensured the fixed number of electrodes: After an initial optimization, including the complete electrode layout, the six electrodes with the maximal current weighting were selected. The applied currents were balanced, while separately preserving the optimized current weightings for anodes and cathodes, respectively. Results from pilot simulations showed reasonable electrode montages and current densities, compared to the original ADMM. Given a fixed electrode size and input current, the number of electrodes significantly affects the current densities and is furthermore important for combined EEG-tES applications. Therefore, it is important to keep this parameter fixed when comparing stimulation montages. *I*_*Total*_ was scaled to 2 mA.

### 2.5 Quantification and Statistical Analysis of Modeling Results

As a result of the forward modeling within the inverse optimization process, current densities [A/m^2^] are given for each node of the FEM head model. Measures for stimulation intensity and spatial extent of the electric field were computed to evaluate the simulated current densities within the individual FEM head models.

Current densities were corrected for the alignment with the electric field by scaling the vector length (the magnitude of the current density) by the absolute of the dot product of each nodes’ normalized vector direction and the target vector orientation. Thereby, parallelity of the stimulation field with the stimulation target was considered. Nodes in GM and WM were included only. Stimulation intensity was defined as the 0.95 percentile including all nodes within 5 mm distance to the target vector. Spatial extent of the electric field was defined, similar to the ‘half-max-radius’, described by Dmochowski and colleagues (2011). The 0.95-percentile was computed as a function of the Euclidean distance to the stimulation target vector for all FEM nodes within a distance to the target vector in an interval from 0 to 100 mm, in steps of 5 mm. The resulting function was interpolated and individually normalized in order to quantify spatial extent independent of stimulation intensity. Spatial extent was quantified as the distance (in mm) at 50% area under the curve of this function. As can be seen in Fig. 2B, especially with respect to the standard montage, the highest intensity is not necessarily centered to the stimulation target. The applied method takes these deviations into account and therefore yields a reliable quantification of stimulation focality. The spatial extent of the electric field reflects the focality of the electric field. Low spatial extent values reflect high focality and vice versa.

**Fig. 2.**
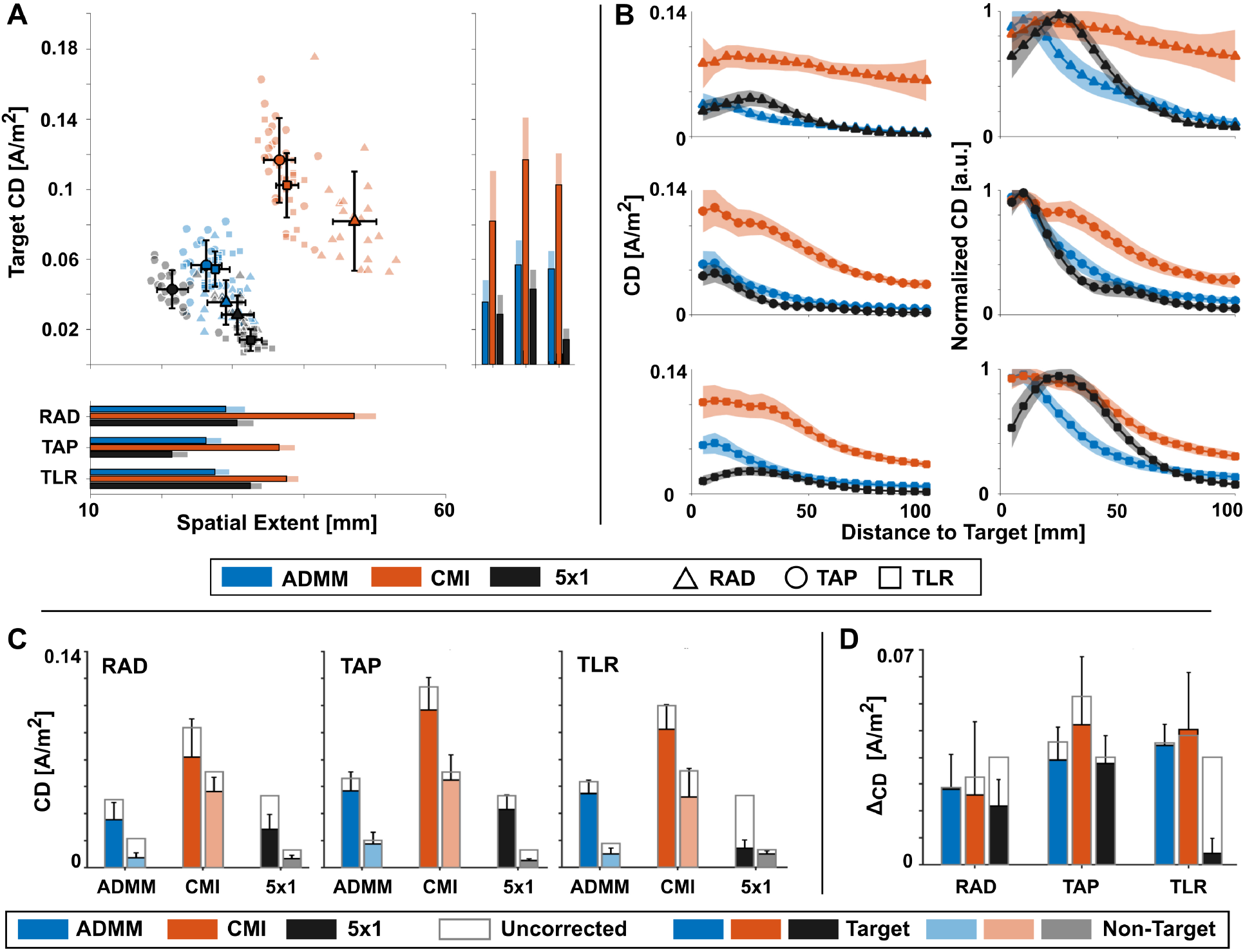
A) Target current densities (CD) plotted against the spatial extent for every method and target orientation. Group averages are represented by colored symbols with black edges (± standard deviation; SD). Single-subjects are plotted in light colors. Bar graphs illustrate the descriptive average target CD and spatial extent, respectively. Light colored stacked bars represent the SD. B) Line plots represent current densities as a function of distance to the target vector (left) for each of the applied methods and target orientations (from top to bottom: radial, tangential_a-p_, tangential_l-r_). Normalized current densities (right) as a function of distance to the target vector illustrate the shape of the electric field, independently of the intensity. C) Descriptive current densities for the target right SPL (solid colored bars) and the non-target left SPL (light colored bars). D) Δ_current densities_ for target (right SPL) and non-target (left SPL) region. Due to the high reliability of the data in A)-C), statistical analysis was only applied to the data shown in D). Results are presented in Tab. 1-3. Corrected current densities are displayed in A) - D). C) and D) additionally show the respective uncorrected current densities as stacked uncolored bars. Error bars and patches represent the sample standard deviations.

In order to compare the specificity of the approaches, the target current density in the right SPL was compared to the left SPL non-target current density (Fig. 2C-D). Non-target current density was computed with respect to the left SPL as described above for the target right SPL. For descriptive reasons, absolute current densities are given for both targets and non-targets in Fig. 2C. Furthermore, values are displayed showing the uncorrected intensity values. Absolute differences between target and non-target current densities (Δ_current densities_) were computed for each method and target orientation. Repeated measures 3×3-ANOVAs were computed to compare the effect of three different stimulation methods (ADMM, CMI, 5×1) and three different target orientations (radial, tangential_a-p_, tangential_l-r_) for stimulation target intensity and focality, as well as hemispheric current density differences, separately. Greenhouse-Geisser correction was applied, if the sphericity assumption was violated. Follow-up paired samples t-tests were computed and *p*-values were corrected for multiple comparisons using the Bonferroni-Holm method (Holm, 1979).

**Tab. 1.** Results of the repeated measures ANOVA testing differences in the Δ_current densities_ of target and non-target. Statistical comparisons were computed across methods (Method: ADMM, CMI, 5×1) and target orientations (Target: radial, tangential_a-p_, tangential_l-r_). Significant main effects for both factors and a first order interaction effect were revealed. *** indicate = *p* < .0001.

| | *p* | *F<sub>df</sub>* | *df* | $\eta_p^2$ |
| --- | --- | --- | --- | --- |
| **Method** | < .0001 | 40.14 *** | 1.3, 26.3 | .67 |
| **Target** | < .0001 | 13.91 *** | 2, 40 | .41 |
| **Method x Target** | < .0001 | 25.02 *** | 2.6, 51.3 | .56 |

Non-parametric Spearman correlations were computed to describe the relationship between intensity and spatial extent for each method and target direction, separately.

Sample means and standard deviations were reported, if not indicated otherwise. Significance level was set to α = 0.05 and effect sizes 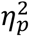 and *r* were reported, respectively. Corrected *p*-values are reported if not indicated otherwise. Detailed information on statistics is provided in Tab. 1-3.

## 3. Results and Discussion

We simulated current densities induced by individually optimized tES montages in realistic 6C FEM head models. Head models were based on an automated segmentation of MRI data for experimental sample sizes. A standard montage was compared with two targeted stimulation montages focusing on stimulation intensity and focality, respectively. Unilateral targets with three different vector orientations were located in the right SPL. We show that individual optimization allows to control variability due to individual structure, as well as target location and orientation, in a sample with the size of typical tES experiments. The results reveal that the choice of the algorithm depends on the specific research question.

### 3.1 Individual Targeting Increases Control over tES-Induced Electric Fields

Electrode montages provided by the ADMM and CMI algorithms showed the expected variation across subjects (see Fig. 3). The 5×1 standard montage was not adapted to the target orientation and individual anatomy, per definition. Thus, electrode positions in the standard montage did not vary across subjects and orientations. For radial targets, ADMM montages showed a pattern comparable to a narrow standard montage with one or more central electrode/s, surrounded by antagonistic electrodes in the close vicinity. By recruiting more than one central stimulation electrode, the ADMM flexibly corrects the current flow, if the radial targets did not project directly on a stimulation electrode at the scalp surface. CMI-derived montages showed one electrode cluster radial to the stimulation target and antagonistic electrode clusters positioned at distributed locations over frontal and temporal regions contralaterally. For tangential targets, ADMM provided two antagonistic electrode clusters that were placed close to one another along the target direction, that is anterior and posterior for tangential_a-p_ and left and right for tangential_l-r_. The spatial extent of the electric field was restricted and non-target brain regions were spared due to the close distance between stimulation electrodes. For the CMI algorithm and the tangential target orientations, two similar electrode clusters were located with increased distance to one another, compared to the ADMM montages. In general, both optimization approaches made extensive use of the high-density electrode layout in order to address the requirements of individual anatomy and variations in target location and orientation. The present data supports the conclusion that the utilization of high-density electrode layouts for the individual optimization of tES montages (Guler et al., 2016) is beneficial, in order to allow the adaptation to the individual anatomy of the participants and to the stimulation target. As depicted in Fig. 3, all montage-target combinations yielded similar dissociation between the target current densities in right SPL and non-target current densities in left SPL, as long as the specific target orientation remained disregarded.

**Fig. 3.**
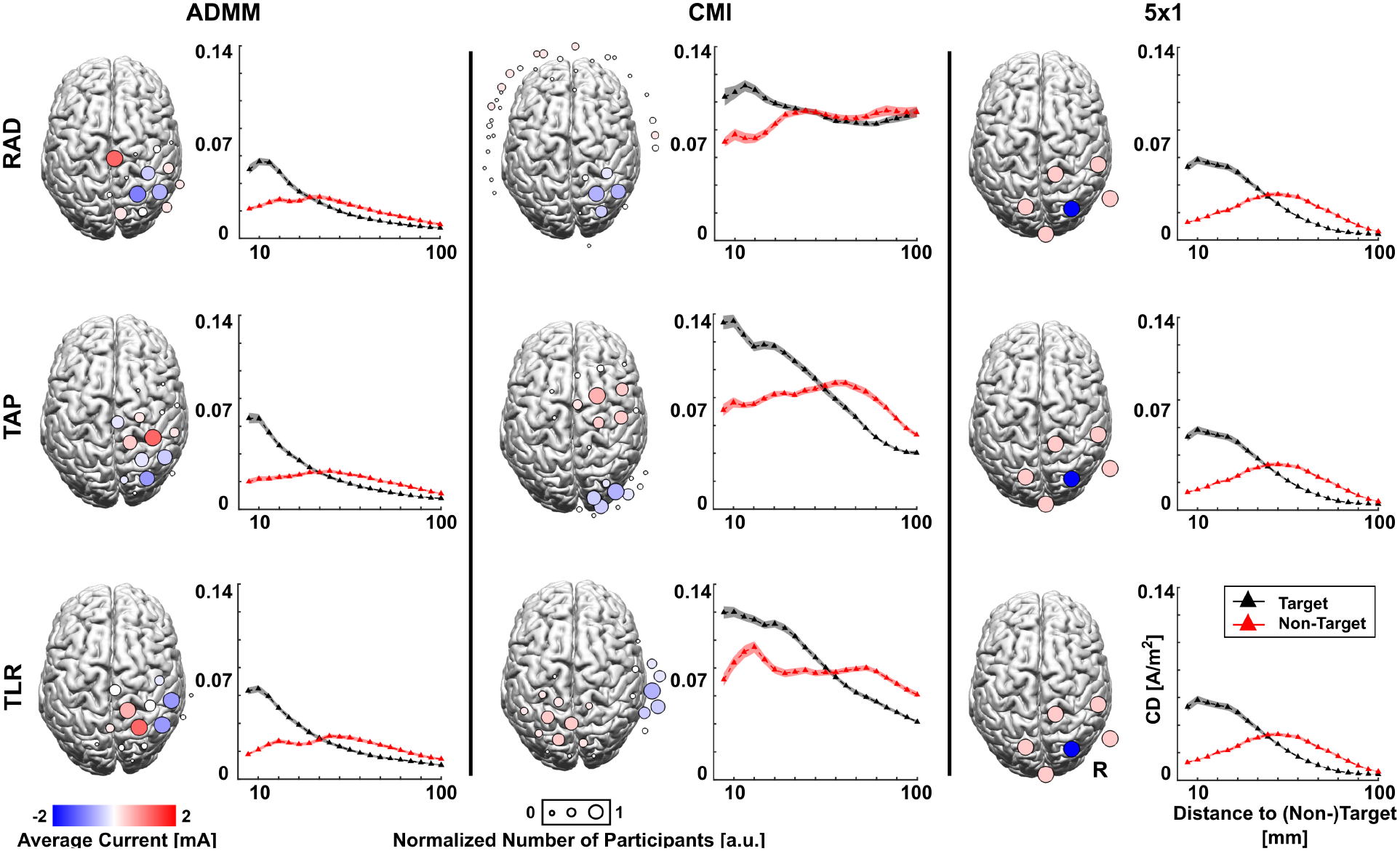
Three-dimensional illustration of the grand average electrode montages that were derived from the focality-optimizing ADMM, intensity-optimizing CMI and a 5×1 standard montage. A template cortical surface (FieldTrip) is plotted, viewed from the top. For each method (from left to right: ADMM, CMI, 5×1) and target orientation (from top to bottom: radial, tangential_a-p_, tangential_l-r_) the electrode sizes reflect the number of participants for whom the respective electrode was part of the montage. Large circles illustrate that for all participants this particular electrode was part of the montage. Smaller circles illustrate fewer usages of the particular electrode. The color of the circles shows the average current applied to the electrode across participants. Red and blue colors indicate that this particular electrode rather had the same polarity across subjects, as well as their relative weighting. White electrodes indicate either small current applied to the electrode or variable electrode polarity across participants. Electrodes that were not part of the resulting electrode montages were omitted, respectively. For each montage illustration, the respective target right SPL (black) and non-target left SPL (red) uncorrected current densities are shown as a function of distance to the target/non-target vector, respectively. Standard 5×1 montages show no variability in the electrode montages and uncorrected current densities, but are depicted here for illustrative reasons.

Exemplary electrode montages for one subject showed similar patterns as described above for the sample (Fig. 4). Current densities resulting from the optimized tES montages were nicely aligned with the target orientation (Fig. 4 and 5), as expected (Dmochowski et al., 2011; Khan et al., 2019; Wagner et al., 2016a). In contrast, the standard montage failed to comply with the orientation of the electric field of the tangential_l-r_ target and the alignment of the electric field with radial and tangential_a-p_ targets was at least suboptimal. To quantify the misalignment of the electric field and the respective target orientation, current densities that were corrected for the orientation of target are reported. Irrespective of the vector field orientation, the current densities were increased in the gyral crowns on a descriptive level (Fig. 5). Similar electric field maxima were previously described as *hotspots* (Datta et al., 2009; Laakso et al., 2015; Opitz et al., 2015).

**Fig. 4.**
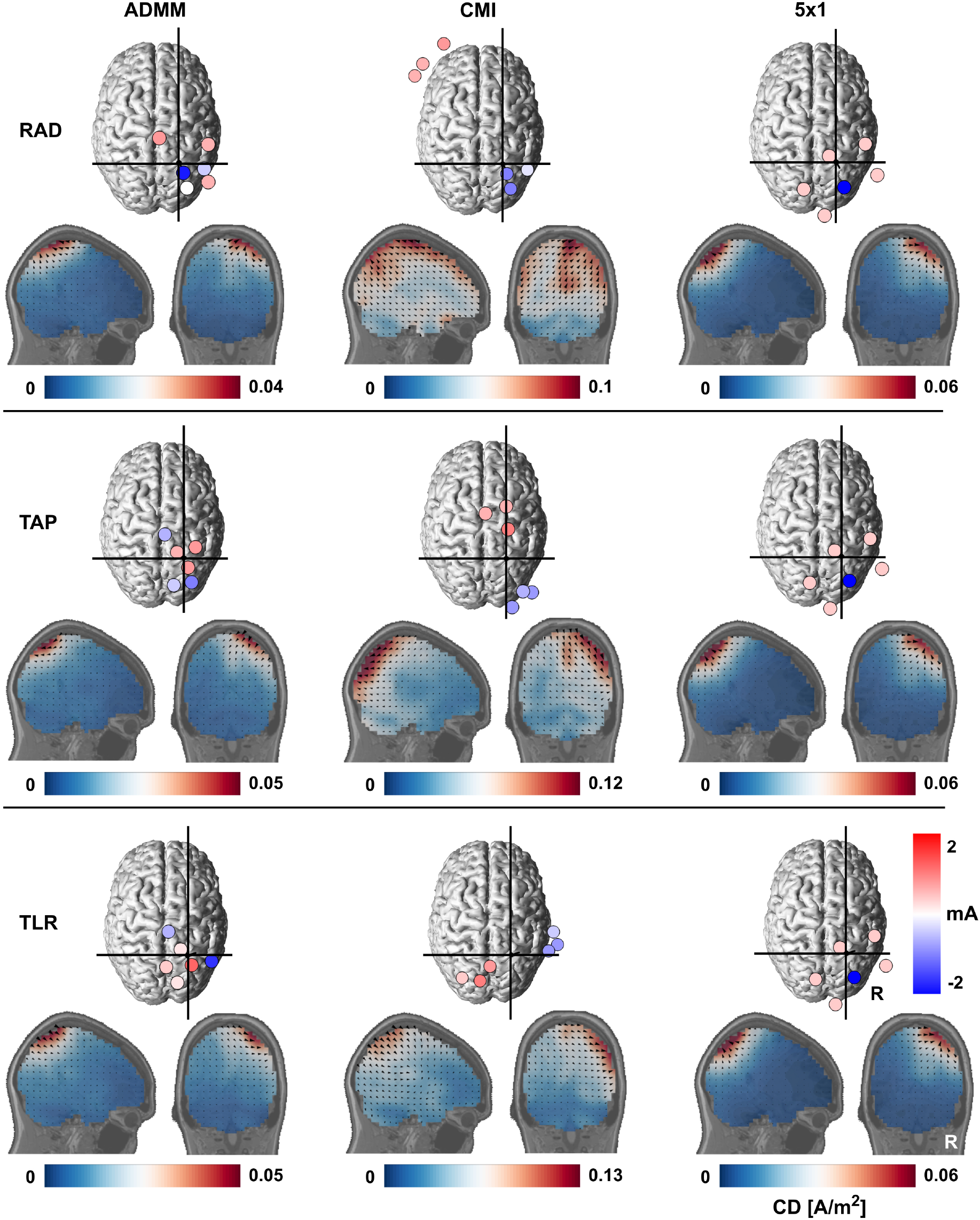
Three-dimensional illustration of electrode montages that were derived from the focality-optimizing ADMM, intensity-optimizing CMI and a 5×1 standard montage for one exemplary participant. The individual cortical isosurface is plotted, viewed from the top. For each method (from left to right: ADMM, CMI, 5×1) and target orientation (from top to bottom: radial, tangential_a-p_, tangential_l-r_) the electrodes are depicted by circles of the same size. The color of the circles reflects the current applied to the respective electrode. Sagittal and coronal slices through the target vector are presented for each electrode montage at the target location. Uncorrected intensities were interpolated on an individual 5 mm grid including GM, WM and CSF and were plotted on top of the individual T1 image. To illustrate the electric field distribution across the brain, custom scales were chosen for each method and target orientation. Three-dimensional vector fields were projected to the respective two-dimensional plane, scaled by the uncorrected current density and plotted on a sparse grid. Vector fields reflect the alignment of targeted tES montages to the target vector orientation (see Fig. 5). Standard 5×1 montages show no variability in the electrode montages and uncorrected current densities, but are depicted here for illustrative reasons.

**Fig. 5.**
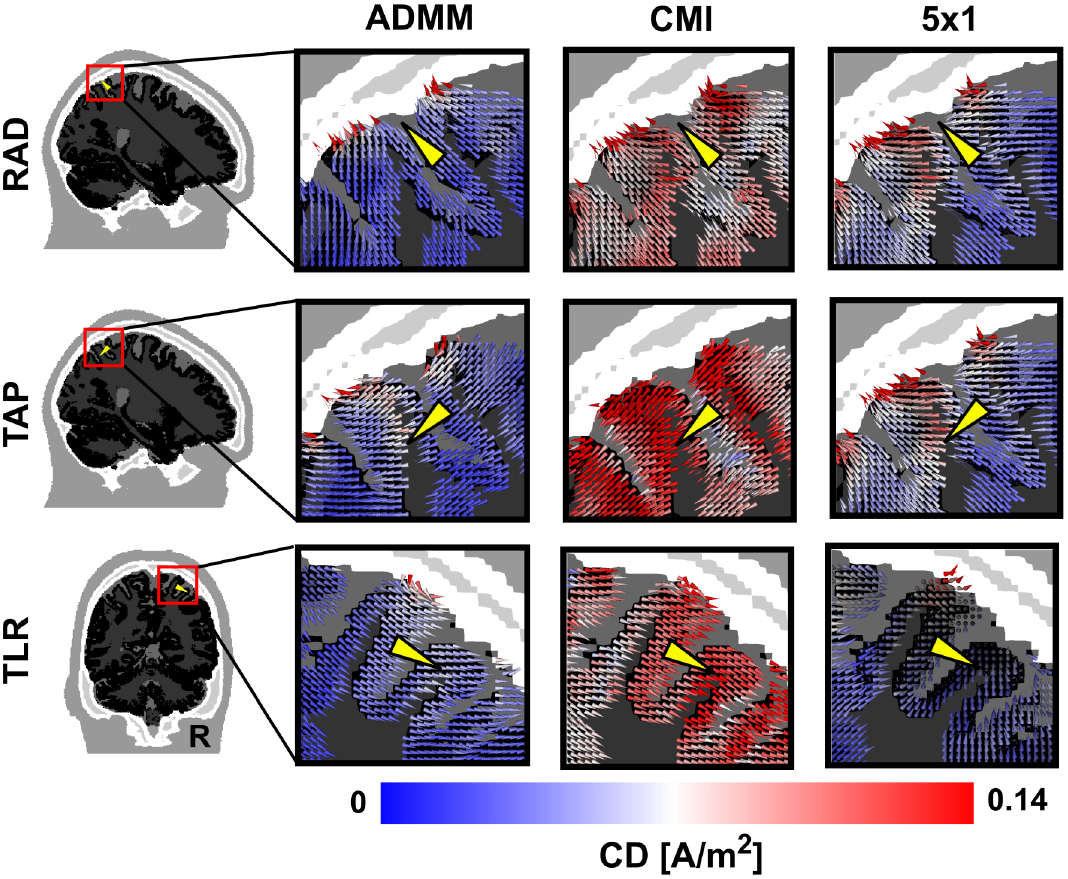
Close-up of the finite-element vector field and target vectors for all methods (from left to right: ADMM, CMI, 5×1) and target orientations (from top to bottom: radial, tangential_a-p_, tangential_l-r_) in one exemplary participant. To optimally depict the field alignment with the target vector orientation, sagittal slices are presented for the radial and tangential_a-p_ targets and a coronal slice was chosen for the tangential_l-r_ target. Vector fields were nicely aligned for both ADMM and CMI with varying spatial extent of the uncorrected current densities and varying maximal current densities. Standard 5×1 montages showed no variability in the electric field, but clear misalignments of the target vector and the electric field were obvious, compared to the targeted electric fields and especially for the tangential_l-r_ target orientation.

Across the three target orientations, corrected target current densities were increased for both ADMM and CMI montages compared to the standard 5×1 montage (Fig. 2; see Supplementary Material). ADMM-optimized electric fields were characterized by small spatial extent, i.e. maximal current densities in close vicinity to the target and steep slopes of current density as a function of distance to the stimulation target (Fig. 2A-B). In contrast, target current densities were highest for the CMI montages for all three target orientations. At the same time, CMI current densities showed the highest spatial extent. Electric fields of the 5×1 standard montages showed varying spatial extent across the three target orientations. Current densities of the 5×1 montages showed a strong bias that peaks at around 20 mm distance to the stimulation target for radial and tangential_l-r_ orientations (Fig. 2B).

The difference between target and non-target current densities (Δ_current densities_) revealed a significant interaction between method and target orientation (Fig. 2D, Tab. 1). Across methods, Δ_current densities_ for radial target orientation were comparable. Surprisingly, follow-up t-tests confirmed significantly increased Δ_current density_ for the tangential_a-p_ target orientation using CMI compared to ADMM and 5×1 montages, although CMI resulted in more distributed electric fields (see Fig. 2 and 3). The present simulations indicate that CMI is capable of defining electrode montages which can sufficiently dissociate electric field strength between non-target (left SPL) and target (right SPL) regions.

Both ADMM and CMI showed higher Δ_current densities_, compared to 5×1 for the tangentiall-r target (Tab. 2). Within the standard 5×1 montage, significant Δ_current densities_ were observed across all three target orientations (Tab. 3). In sum, these results reflect the missing adaptation of electric fields derived from the 5×1 montage to individual anatomical properties and target orientations. In contrast, both ADMM and CMI algorithms were able to adapt to different target orientations and still result in relatively high current densities.

**Tab. 2.**
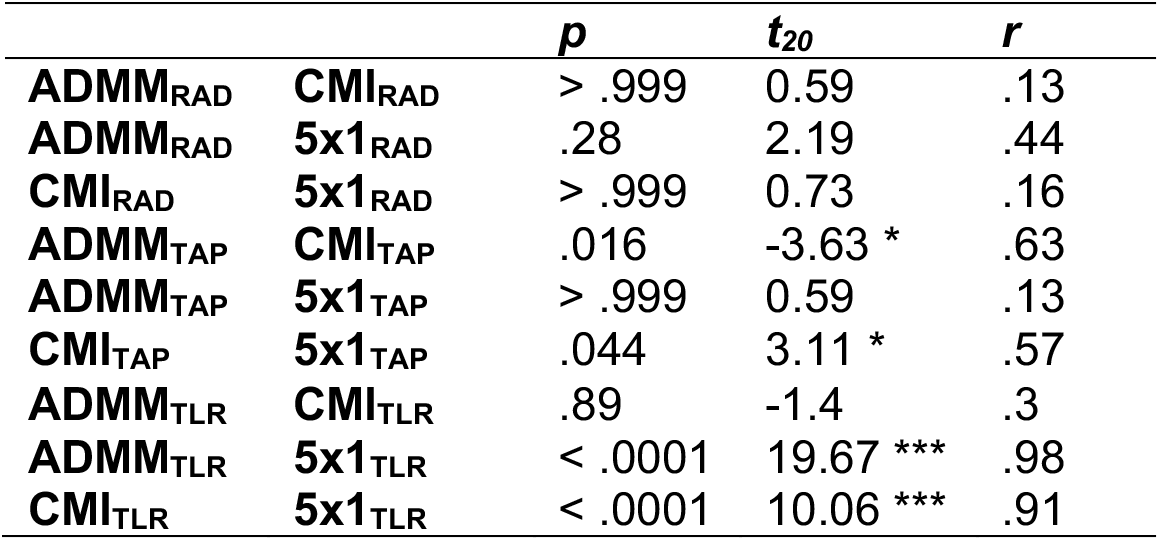
Follow-up t-tests of the Method x Target interaction showing differences in Δ_current densities_ across methods. CMI solutions for the tangential_a-p_ target orientation showed increased Δ_current densities_ compared to ADMM and 5×1. For the tangential_l-r_ orientation, both ADMM and CMI electric fields showed increased Δ_current densities_ compared to the standard 5×1 montage. * indicate = *p* < .05. *** indicate = *p* < .0001.

**Tab. 3.** Follow-up t-tests of the Method x Target interaction showing differences in Δ_current densities_ across target orientations. Except for the tangential_l-r_ orientation in 5×1 montages, for all remaining tangential target orientation-method combinations Δ_current densities_ were increased compared to the radial target. The additional significant difference in Δ_current densities_ between tangential_a-p_ and tangential_l-r_ orientations, as well as the overall effect sizes represent the increased variability of Δ_current densities_ with application of the 5×1 standard montage. * indicate = *p* < .05. ** indicate = *p* < .001. *** indicate = *p* < .0001.

|  |  | *p* | *t*<sub>20</sub> | *r* |
| --- | --- | --- | --- | --- |
| **ADMM<sub>RAD</sub>** | **ADMM<sub>TAP</sub>** | .018 | -3.55 * | .62 |
| **ADMM<sub>RAD</sub>** | **ADMM<sub>TLR</sub>** | < .001 | -5.03 ** | .75 |
| **ADMM<sub>TAP</sub>** | **ADMM<sub>TLR</sub>** | .56 | -1.76 | .37 |
| **CMI<sub>RAD</sub>** | **CMI<sub>TAP</sub>** | .013 | -3.77 * | .64 |
| **CMI<sub>RAD</sub>** | **CMI<sub>TLR</sub>** | .012 | -3.85 * | .65 |
| **CMI<sub>TAP</sub>** | **CMI<sub>TLR</sub>** | > .999 | 0.26 | .06 |
| **5x1<sub>RAD</sub>** | **5x1<sub>TAP</sub>** | < .0001 | -7.77 *** | .87 |
| **5x1<sub>RAD</sub>** | **5x1<sub>TLR</sub>** | < .0001 | 7.38 *** | .86 |
| **5x1<sub>TAP</sub>** | **5x1<sub>TLR</sub>** | < .0001 | 14.09 *** | .95 |

Furthermore, radial targets showed lower current densities, compared to both tangential orientations for ADMM and CMI (Dmochowski et al., 2011). In the present study, stimulation targets were located in intermediate sulcal depth for all target orientations. Previous simulations of targeted tES investigated the effect of target depth, comparing superficial radial targets with intermediate or deep tangential targets (Dmochowski et al., 2011; Wagner et al., 2016a). Herein, current densities were reduced, with increasing depth of the stimulation target (Wagner et al., 2016a). However, these studies were inconclusive on how targeted stimulation can cope with varying target orientations, independent of target depth. We conclude that - in intermediate depth - targets with a tangential orientation can be stimulated with a relatively higher current density, compared to radial targets. Nevertheless, the same does not necessarily hold for stimulation targets in more superficial cortical areas, but has to be investigated in future studies.

As described previously (Dmochowski et al., 2011), we observed a trade-off between intensity and focality, when comparing focality-optimizing (ADMM) and intensity-optimizing (CMI) algorithms for targeted tES. Although this trade-off was reported across methods, the inverse relationship between intensity and focality was apparent across subjects, in the present data (CMI_TLR_: ρ = −0.58, *p* = .041; 5×1_RAD_: ρ = −0.78, *p* < .001; 5×1_TAP_: ρ = −0.64, *p* = .014; CMI_TAP_: ρ = −0.52, *p* = .096, (*p*_uncorrected_ = .016)). No such relationship was observed for electric fields derived from ADMM for any of the three target orientations. Taken together, some subjects showed a systematic increase in stimulation intensity *and* focality for some CMI and 5×1 montages - relative to other subjects. Critically, in case of ADMM, the individual optimization can compensate for individual variation, making the stimulation more reliable across subjects, compared to CMI and 5×1, at least with respect to the spatial extent of the electric field.

In previous simulation studies, electric field hotspots in gyral crowns and sulcal pits (Datta et al., 2009) were described repeatedly. Hotspots depend on anatomical properties like CSF thickness and sulcal depth (Opitz et al., 2015) and vary across subjects (Laakso et al., 2015). Using targeted tES, it might be possible to minimize hotspots outside the target, or make use of them in case they match the stimulation target. Furthermore, the field orientation might play a critical role in the handling of hotspots, and it was mentioned before that the electric field orientation can partly be controlled by using the ADMM and CMI algorithms to optimize stimulation montages. As illustrated in Fig. 5, and following the mathematical maximum principle (Wagner et al., 2016a), the highest current densities can be observed in the gyral crowns. At the same time, for both the ADMM and CMI algorithms, electric fields are aligned to the respective target orientations. Critically, for those targets oriented tangentially to the scalp surface, the field is also oriented tangentially with respect to the scalp surface. This was also observed for the electric field at the gyral crowns in the close vicinity to the target. Consequently, the electric field in the gyral crowns would be orthogonal to the orientation of cortical pyramidal cells in these areas, affecting the subthreshold modulation of the cells membrane potential (Radman et al., 2009). Thus, in the case of a tangential field orientation the electric field would probably have only limited physiological effect on pyramidal cells in the gyral crowns.

Besides the control over the field orientation, the target current density was optimized, while simply ignoring the electric field in non-target regions in the case of CMI (Fig. 2). This is intended to improve the probability that individual electric fields will take physiological effect within the target (Antal and Herrmann, 2016; Reato et al., 2013). In contrast to the expectations, both the ADMM and the CMI were not able to homogenize the variability of target intensities across subjects. Instead, the individually optimal target intensity (under the given constraints) was computed, which seems to be limited by the individual anatomical properties of the head model. Rather, ADMM and CMI both exhibited their effect by orienting the electric field (peaks) towards the stimulation target and herein were less prone to varying anatomy, target location and orientation. Taken together, both optimization algorithms were able to reliably reduce the bias of electric field maxima, with respect to the stimulation target. Due to the varying target intensities, the ad-hoc estimation or post-hoc evaluation of individual tES-induced target current densities still remains highly recommended in order to evaluate effects of individual anatomy on the behavioral or neurophysiological efficacy of tES.

Furthermore, other than the applied algorithms for targeted tES are available (Dmochowski et al., 2017, 2011; Guler et al., 2016; Ruffini et al., 2014). In principal, these algorithms intend to optimize the target current density compared to standard montages. However, specific applications demand different algorithm parameters. Depending on the needs of the respective application, it is possible to control the spatial extent of the electric fields, maximize the target current densities or adapt the electric field orientations with respect to predefined stimulation targets.

In sum, we conclude that targeted tES allows some level of control over the electric field with respect to individual anatomical properties. For the first time, evidence was provided for the application of targeted tES in order to increase control over individual variation of target location and orientation in a sample sufficiently sized for experimental application. The focality-optimizing ADMM yields the most reliable results across stimulation target orientations, compared to CMI and the 5×1 montages. In line with previous results, CMI is well suited to maximize the target current density, especially for tangential_a-p_ targets (cf. Dmochowski et al., 2011; Khan et al., 2019). Interestingly, Δ_current densities_ were preserved and partly even increased - compared to ADMM and standard 5×1 montages - for the evaluated right SPL target and left SPL non-target regions. In real applications of tES, the probability that the 5×1 montage is able to deliver a sufficient electric field in the target is highly dependent on the individual anatomy and the target location and orientation. Since we chose a rather broadly distributed standard montage, we might have even underestimated the bias that is introduced by a standard montage. The bias most likely would be increased, if the surrounding electrodes were placed closer to the center electrode. Furthermore, the applied spherical quantification of intensity included areas in the vicinity of 1 cm around the target vector. As can be seen in Fig. 4, this might have further overestimated the efficacy of the 5×1 montage for all three target orientations, especially when corrected for target orientation.

Taken together, the present study design enabled the individual simulation of electric fields in the head models of twenty-one subjects. A high-density electrode layout was used to allow the ADMM and CMI to flexibly adapt stimulation montages to variations of individual structure, as well as target location and orientation. Based on the presented data, we follow that different algorithms for targeted tES are able to increase the control over specific electric field properties, with respect to the target current density, the orientation, or the spatial extent of the electric field.

### 3.2 The Role of Targeted tES for Physiologically Effective Stimulation

The estimation of intracranial electric fields is necessary in order to understand the physiological mechanisms that are affected by tES. FEM modeling can provide information that is required in order to close the gap between transcranial network stimulation in humans and intracranial stimulation in animals on meso- and microscopic level (Bestmann et al., 2015; Reato et al., 2013). In the present study target current densities ranged from 0.007 A/m^2^ (5×1, tangential_l-r_) to 0.176 A/m^2^ (CMI, radial). These observed target intensities fit into the range of previously reported FEM simulation results (Tab. 4) (Boayue et al., 2018; Datta et al., 2012; Dmochowski et al., 2011; Opitz et al., 2015; Wagner et al., 2016a, 2014).

**Tab. 4.** Minimum and maximum target current densities [A/m^2^] across subjects are depicted for every method and target orientation. In order to directly compare the present results with previous FEM simulation studies, electric field strength [V/m] was computed from current densities, with GM conductivity of 0.33 S/m. Spatial extent of the electric fields is given in [mm].

|  | A/m<sup>2</sup> |  | V/m |  | mm |  |
| --- | --- | --- | --- | --- | --- | --- |
|  | min | max | min | max | min | max |
| **ADMM<sub>RAD</sub>** | .019 | .074 | .057 | .224 | 26 | 33 |
| **ADMM<sub>TAP</sub>** | .036 | .082 | .109 | .248 | 23 | 29 |
| **ADMM<sub>TLR</sub>** | .042 | .075 | .127 | .227 | 23 | 31 |
| **CMI<sub>RAD</sub>** | .053 | .176 | .161 | .533 | 42 | 53 |
| **CMI<sub>TAP</sub>** | .066 | .163 | .2 | .494 | 34 | 42 |
| **CMI<sub>TLR</sub>** | .069 | .14 | .209 | .424 | 34 | 40 |
| **5x1<sub>RAD</sub>** | .02 | .051 | .061 | .155 | 25 | 34 |
| **5x1<sub>TAP</sub>** | .019 | .063 | .058 | .191 | 19 | 28 |
| **5x1<sub>TLR</sub>** | .007 | .035 | .021 | .106 | 29 | 36 |

In previous studies, *in vitro* and *in vivo* recordings reported subthreshold modulation of neuronal activity that were induced by electric fields with peak intensities at 0.2 - 0.5 V/m (approx. 0.066 - 0.165 A/m^2^) using alternating current stimulation at several frequencies (Deans et al., 2007; Francis et al., 2003; Fröhlich and McCormick, 2010). Nevertheless, the mechanism that drives those effects still is not resolved. Proposed explanations range from slight modulations of neural activity, like stochastic resonance to the full entrainment of neural activity by the electric field, depending on the applied stimulation amplitude, frequency and the relation between the exogenous electric field and endogenous neural activity (Liu et al., 2018; Reato et al., 2013). Due to the complexity of the problem, the comparability between transcranial stimulation in humans and intracranial stimulation in (anesthetized) animals or in vitro recordings is limited. Due to varying tissue conductivities (Aydin et al., 2017, 2014; Huang et al., 2017), effects of network electric activity (Francis et al., 2003; see Reato et al., 2013) and the effective state of the stimulated neuronal population in the behaving human participant (Alagapan et al., 2016; Neuling et al., 2013), already relatively weak electric fields might potentially modulate neuronal activity in specific cases, compared to isolated neurons or neural populations. Still, in line with previous modeling results (Datta et al., 2012; Dmochowski et al., 2011; Opitz et al., 2015; Wagner et al., 2016a, 2014) the present results indicate that neural modulation, in principle is possible when applying tES in human participants. Nevertheless, tES is acting at the lower end at which electric fields were reported to modulate neural activity (see Fig. 2) (Deans et al., 2007; Francis et al., 2003; Fröhlich and McCormick, 2010; Reato et al., 2013).

Recently, a method to measure intracranial electromagnetic fields using MRI and tDCS was described. This non-invasive method allows to directly validate the simulated electric field, based on human in vivo measurements. First results indicate that FEM models yield a reasonable approximation to the measured electric fields (Göksu et al., 2018). Additionally, estimated electric fields from specific FEM head models showed a good correspondence when compared to intracranial recordings from human patients (Huang et al., 2017; Lafon et al., 2017) and non-human primates (Kar et al., 2017; Opitz et al., 2016). The observed modeling errors in these studies were mostly attributed to tissue conductivities (Göksu et al., 2018; Huang et al., 2017). In the large body of FEM applications these are not adapted to individually varying conductivity values and thus might lead to errors in the estimation of intracranial current densities induced by tES (Aydin et al., 2014; Schmidt et al., 2015). At the same time it is not yet possible to estimate conductivities non-invasively and for individual FEM head models, except for some specific applications (Arumugam et al., 2018; Aydin et al., 2014; Fernandez-Corazza et al., 2018). Therefore, the inclusion of individual conductivities remains an important future refinement of FEM head models. Nonetheless, anatomical features other than tissue conductivities introduce variability to electric fields across individuals. Among these, CSF distribution, sulcus morphology, as well as the depth of the stimulation target were shown to have the greatest impact on electric field simulations (Laakso et al., 2015; Opitz et al., 2015). The FEM approach applied here specifically aims at the spatial arrangement of these tissue compartments inside the head and is able to accurately model these features (Göksu et al., 2018; Huang et al., 2017). Consequently, individual targeting of tES induced electric fields increases the level of control over the same individual features.

When combining FEM modeling with real tES applications, the estimated fields might provide an a priori estimation, or a posteriori explanation of the behavioral or neurophysiological efficacy of tES, even for standard montages (e.g. Kasten et al., 2019). It has to be noted that the input current to the stimulation electrodes is limited by safety constraints and experimental considerations (e.g. minimizing skin sensations). By individually optimizing the electric field, the full potential of the applied field can be shaped with respect to the stimulation target location and orientation in order to increase the efficacy of tES in real applications, compared to standard montages.

### 3.3 Automatic 6C Segmentation for Individual FEM Modeling in Experimental Samples

In order to understand the subtle and complex consequences of tES, an integration of simulations of the individual electric fields into experimental applications of tES is indispensable. At the same time, realistic FEM head models allow a reasonable estimate of the electric field distribution, but are effortful in terms of computational demands and expert resources (Huang et al., 2013). A new MATLAB-based segmentation tool was implemented to segment the T1 and T2 MRI data into six compartments in an automated and standardized way (https://doi.org/10.5281/zenodo.3365193). The applied algorithm reliably produced specific and anatomically reasonable results, as presented in Fig 1D. Segmentation results were validated by careful visual inspection of the segmentation for every subject, and comparing it to the initial T1 and T2 MRI images. Furthermore, no significant leakage artifacts were revealed by the modeling results of current flow (e.g. Engwer et al., 2017; Vorwerk et al., 2017), speaking in favor of the segmentation accuracy.

However, like every head model this one also represents an approximation to the real anatomy of the head. Systematic segmentation bias was predominantly observed in fronto-basal regions. For example, a systematic bias was introduced by the supine position of the participants in the MR scanner during data acquisition. This position provokes the brain to move towards the cerebral fossa of the occipital bone, thereby displacing CSF to more anterior regions (Rice et al., 2013). Furthermore, in orbital and basal regions, skull openings like the optic canal and the foramen magnum might have been overestimated by the segmentation algorithm. Since modeling of skull openings has a substantial effect on modeling of current flow (Datta et al., 2010; Lanfer et al., 2012; Mekonnen et al., 2012; Oostenveld and Oostendorp, 2002), we applied a segmentation based on the integration of T1 and T2 imaging data to increase the accuracy of bone estimates (Aydin et al., 2014; Wagner et al., 2014). The largest effects of skull openings are expected for sources and electrodes close to the opening (Lanfer et al., 2012; Mekonnen et al., 2012; Oostenveld and Oostendorp, 2002). Thus, effects of fronto-basal segmentation inaccuracies were expected to be small with respect to the applied posterior stimulation targets in the current study. A reduced head model was computed focusing on the reliability of the segmentation, neglecting for example meninges as well as the eye and sub-compartments of the skin (cf. Laakso et al., 2015). Since no air compartment was included in the model, air-filled sinuses were misclassified, overestimating the conductivity for the respective voxels in frontal skull regions.

The segmentation applied here extends the application-oriented family of automatic segmentation tools. In contrast to other available automated segmentation tools (Huang et al., 2013; Windhoff et al., 2013), BONE_S_ was included in the model. Due to the location of stimulation targets in the current study, electrodes were predominantly placed over parietal regions. Here, BONE_S_ affects the current flow, in a comparable (but inverted) fashion like frontal air cavities that were incorporated previously in head models (Huang et al., 2013). Due to higher conductivity of BONE_S_, compared to BONE_C_, the current flow is redirected from an otherwise radial flow through the low-conductive BONE_C_ (Wagner et al., 2014). This effect might lead to an increased outspread of the electric field across the underlying cortical surface and counteracts the insulating effect of BONE_C_ in regions of thick skull (Opitz et al., 2015). Although the effect of BONE_S_ is negligible with respect to the current density in the directly underlying cortex (Opitz et al., 2015) it yields a complex effect on the overall cortical current density and on the orientation of the current (Opitz et al., 2015; Wagner et al., 2014).

Besides the indicated limitations, the resulting simulations support the view that the head model was sufficiently suited for the simulation of electric fields. Importantly, in contrast to more specific but also more effortful semi-automatic segmentation methods, which require manual corrections, automatic algorithms are able to speed up and standardize the segmentation of MRI data. These features are especially beneficial for the broad application of FEM modeling in clinical and experimental settings with larger sample sizes. At the same time, the modeling of current flow as a standard application for tES might be beneficial to promote the individualized application of tES and to extend the understanding of its’ underlying mechanisms.

## 4. Conclusion

Individual anatomical properties lead to individual variability in the potential for tES taking effect. Critically, targeting of tES electric fields is able to increase the level of control over the individual intracranial current densities, compared to less accurate and less reliable standard montages. The focality-optimizing ADMM algorithm allows a balance between spatial extent and target intensity, while flexibly adapting to target orientation and individual anatomy. The intensity-optimizing CMI algorithm improves the target intensity of the electric fields estimated in individual head models, thereby raising the overall chance of a physiological tES effect. Furthermore, current densities resulting from unilateral stimulation were described, showing a reasonable dissociation between target and contralateral non-target regions, even for the intensity-optimizing CMI approach. Finally, the experimental sample size allowed insight into the stimulation intensity-focality relation across subjects. Interestingly, in some cases, this relation was opposing the previously described trade-off of stimulation intensity and focality across stimulation montages.

In conclusion, FEM simulation results might help to increase the physiological interpretability of tES effects using targeted tES. We propose that a priori definition of individual stimulation montages and the detailed analysis of estimated electric fields has potential to improve the understanding of mechanisms underlying tES and thus its’ effectiveness in future applications.

