## Supplementary Material for "Simulating individually targeted transcranial electric stimulation for experimental application"

Due to the nature of volume conduction, in many applications of tES the applied electric field indirectly stimulates brain regions that were not specifically targeted. Whereas the above-described quantification addresses the question whether a specific stimulation montage is able to reasonably stimulate the target, another important question is, whether other stimulated brain regions may have given rise to a tES effect. Thus, we computed the global stimulation intensity of the electric field. Since the target vector direction in any other than the target region is not informative at this point, the intensity was not corrected for parallelity to the stimulation target. Using a gaussian kernel, current density forward solutions were interpolated on the same individual 5 mm grids that were utilized for target definition. Each grid-point was labeled according to the respective AAL-region in MNI-space and region-wise intensities were determined as the 0.95-percentile, respectively.

Results reflect similar effects as revealed by the local quantification of target current densities and spatial extent. The ADMM algorithm showed highest intensities in parietal cortex, and some intensity in occipital regions. Intensities were limited to the right stimulation target hemisphere, relatively to the left non-target hemisphere. Across all brain regions, CMI showed higher intensities, compared to ADMM and 5x1, but a good dissociation between hemispheres in target parietal areas and occipital areas. In line with the local quantifications, highest current densities in the target hemisphere were observed for the $\text{tangential}_{a\text{-}p}$ target orientation for the CMI algorithm, peaking at parietal and occipital regions. For the standard 5x1 montage, a bias of the electric field was revealed showing highest current densities in occipital regions. Comparable current densities were observed in the target parietal cortex in the right hemisphere.

In tES applications, both aspects a) the electric field showing a reasonable intensity in the stimulation target, while b) being constrained or unrelated to behavioral or neurophysiological tES outcome are critical. A global quantification of the current densities, based on standard brain atlases might contribute to the understanding the neurophysiological effector of tES outcome by equally considering a) and b).

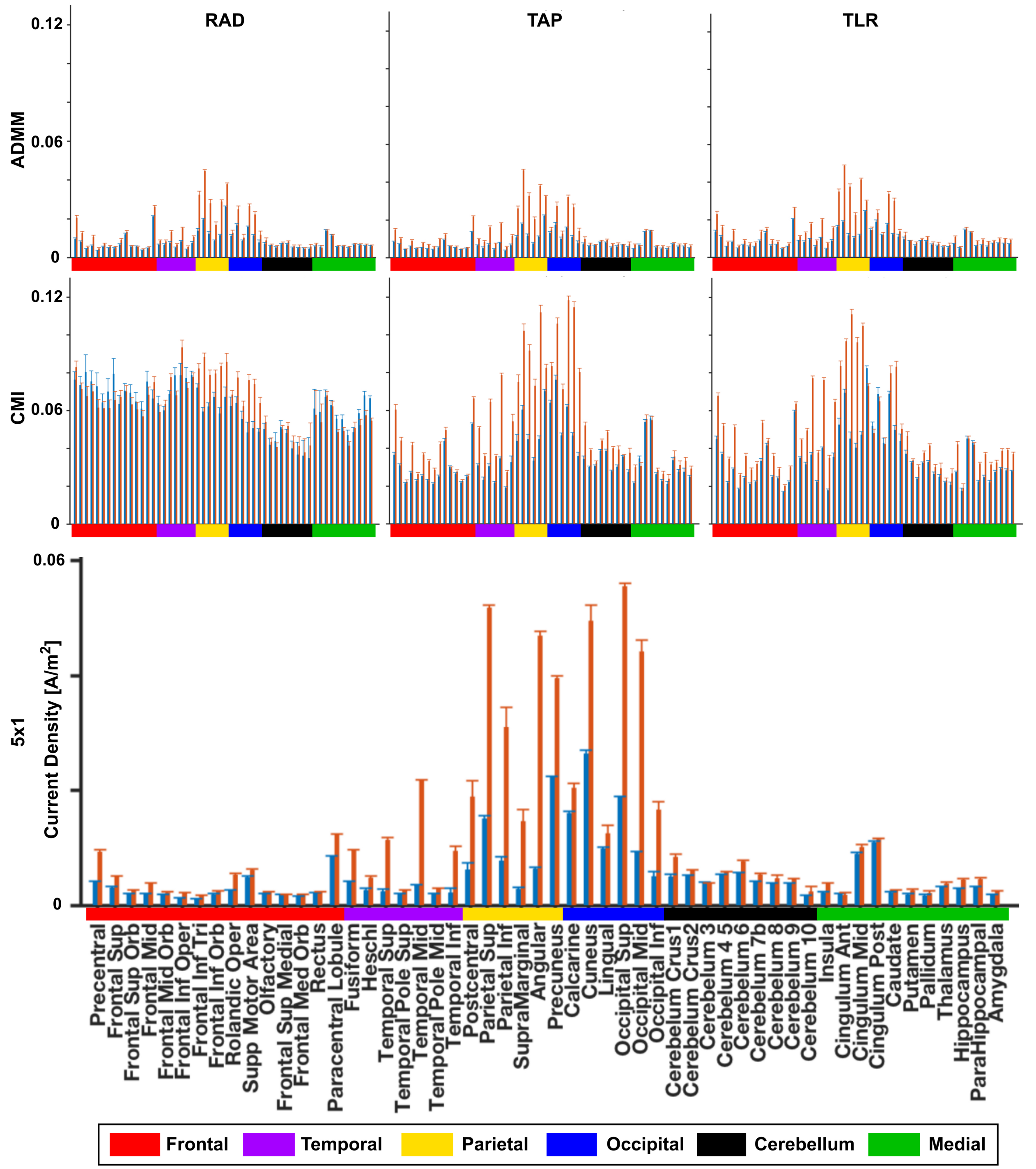


**Suppl. Fig. 1.** 95%-percentile current densities (sample mean ± SEM) are shown for all of 108 bilateral AAL-regions. Bilateral regions were clustered and plotted as red (right, target hemisphere) or blue (left, non-target hemisphere) bars, respectively. Results are depicted for each of three target orientations (radial, RAD; tangential$_{a\text{-}p}$, TAP; tangential$_{l\text{-}r}$, TLR) and methods (ADMM, CMI, 5x1). No differences across target orientations was present for the standard 5x1 induced current densities, therefore only one plot is shown in this figure. Names of the regions are depicted exemplary for the standard 5x1 montage and color-labeled for coarser brain regions. Color-labels in the remaining plots refer to the same regions.
